# Evidence for the priming effect in a planktonic estuarine microbial community

**DOI:** 10.1101/030916

**Authors:** Andrew D. Steen, Lauren N. M. Quigley, Alison Buchan

**Affiliations:** Department of Earth and Planetary Sciences University of Tennessee, Knoxville, USA; Department of Microbiology University of Tennessee, Knoxville, USA

## Abstract

The ‘priming effect’, in which addition of labile substances changes the rem-ineralization rate of recalcitrant organic matter, has been intensively studied in soils, but is less well-documented in aquatic systems. We investigated the extent to which additions of nutrients or labile organic carbon could influence remineralization rates of ^14^C-labeled, microbially-degraded, phytoplankton-derived organic matter (OM) in microcosms inoculated with microbial communities drawn from Groves Creek Estuary in coastal Georgia, USA. We found that amendment with labile protein plus phosphorus increased remineralization rates of degraded, phytoplankton-derived OM by up to 100%, whereas acetate slightly decreased remineralization rates relative to an unamended control. Addition of ammonium and phosphate induced a smaller effect, whereas addition of ammonium alone had no effect. Counterintuitively, alkaline phosphatase activities increased in response to the addition of protein under P-replete conditions, indicating that production of enzymes unrelated to the labile priming compound may be a mechanism for the priming effect. The observed priming effect was transient: after 36 days of incubation roughly the same quantity of organic carbon had been mineralized in all treatments including no-addition controls. This timescale is on the order of the typical hydrologic residence times of well-flushed estuaries suggesting that priming in estuaries has the potential to influence whether OC is remineralized in situ or exported to the coastal ocean.

## 1 Introduction

The ‘priming effect’ refers to changes in the remineralization rate of less bioavailable organic matter (OM) in response to the addition of more bioavailable substances (Kuzyakov et al., 2000; Jenkinson et al., 1985). Although this effect has been the subject of intensive study in soils, it has only recently begun to attract substantial attention in aquatic systems (Guenet *et al*., 2010; Bianchi, 2011; Bianchi *et al*., 2015). Among aquatic systems, the priming effect may be particularly relevant in estuaries, where labile organic matter (OM, for instance autochthonous production) mixes with more recalcitrant OM, such as aged terrestrial OM and recalcitrant marine OM (Guenet *et al*., 2010).

Despite the voluminous evidence for the priming effect in soils (Kuzyakov, 2010), the evidence for priming in aquatic systems is more ambiguous. Several studies using unlabeled labile organic matter to aquatic ecosystems showed by mass balance that additions of labile OM must have stimulated oxidation of more recalcitrant OM (De Haan, 1977; Shimp & Pfaender, 1985; Farjalla *et al*., 2009); other nvestigators in freshwater environments have not found evidence for the priming effect (Bengtsson *et al*., 2014; Catalán *et al*., 2015), while Bianchi *et al*. (2015) observed priming of an estuarine bacterial isolate of Acinetobacter induced by a disaccharide or algal exudate. With the exception of Farjalla *et al*. (2009), which concerns a tropical lagoon, these studies were not performed in estuaries. Perhaps more importantly, the priming effect refers to changes in remineralization of recalcitrant OM in response to the addition of more labile OM and/or nutrients. It can be challenging to distinguish remineralization of labile versus recalcitrant OM using a mass-balance approach, in which only total fluxes of CO_2_ are measured, because these approaches do not distinguish between oxidation of pre-existing, recalcitrant OM and added labile OM.

To assess the extent to which additions of labile OM and/or nutrients may influence the remineralization rates of recalcitrant OM in coastal estuaries, we performed microcosm experiments and monitored the remineralization of degraded, phytoplankton-derived organic matter by a surface water microbial community collected from a temperate coastal estuary (Grove’s Creek, Georgia, USA). Phytoplankton were labeled with ^14^C so fluxes of ^14^CO_2_ derived from phytoplankton-derived OM could unambiguously be distinguished from unlabeled CO_2_ derived from labile carbon. Periodic measurements of cell abundance, extracellular enzyme activities, and dissolved organic matter (DOM) fluorescence provided insight into the mechanisms of interactions between labile OM, nutrients, and phytoplankton -derived OM. These microcosms provided a tractable experimental system in which to assess the influence of simple (acetate) versus complex (protein) labile OM as well as nutrient addition (N or N+P) on degraded OM in estuaries.

## 2 Material & Methods

### 2.1 Generation of ^14^C-labeled organic matter

The marine phytoplankter *Synechococcus* sp. strain CB0101 was grown on SN15 medium (750 mL filtered seawater, 250 mL distilled water, 2.5 mL 3.53 M NaNO_3_, 2.6 mL 352 mM K_2_HPO_4_, 5.6 mL 342 mM Na_2_EDTA · 2H_2_O, 2.6 mL 37.7 mM Na_2_CO_3_, 1 mL 737 μM cobalamin, 1 mL cyano trace metal solution [400 mL distilled water, 100 mL 297 mM citric acid · H_2_O, 100 mL 229 mM ferric ammonium citrate, 100 mL 27 mM MnCl_2_ · 4H_2_O, 100 mL 17.8 mM Na_2_MoO_4_ · 2H_2_O, 100 mL 859 μM Co(NO_3_)_2_ · 6H_2_O, 100 mL 7.7 mM ZnSO_4_ { 7H_2_O]) in a sealed, 4-liter flask in the presence of 0.5 mCi NaH^14^CO3^−^ (MP Biomedicals, Santa Ana, CA) under artificial illumination on a 12-hr/12-hr cycle at 28°C. Stationary phase cultures were collected on 0.22 μM Supor filters (Pall Corporation, Port Washington, NY) and resuspended in artificial seawater (ASW) (Sigma Sea Salts, 20 g/L [Sigma-Aldrich, St. Louis, MO]), pH 8.1. A microbial community inoculum (collected at Bogue Sound, NC, from the dock of the Institute of Marine Sciences, University of North Carolina-Chapel Hill) was added to the phytoplankton biomass at 1% v/v and incubated in the dark at room temperature for 45 days. During the course of the incubation, the quantity of O^14^C was periodically measured using a Perkin-Elmer TriCarb 2910-TR liquid scintillation analyzer (Perkin-Elmer, Waltham, MA). The con-centration of remaining OC was calculated by assuming the specific activity of degraded, phytoplankton-derived OC was equal to the specific activity of DI^14^C in the growth medium.

Phytoplankton-derived OC decay was modeled according to first-order kinetics:

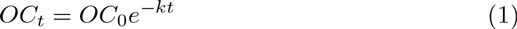

where *OC_t_* is the concentration of organic carbon at time *t, OC*_0_ is the initial concentration of organic carbon, and *k* is the decay rate constant. *k* was determined from a nonlinear least squares regression of the OC concentration data to equation 1, and half-life was calculated as *t*_1/2_ = ln(2)/*k*. At the end of this initial degradation phase, ^14^C-POM was collected by filtration (0.22 μm), resuspended in ASW (salinity = 20) and heat killed by boiling for 5 min. The POM was allowed to return to room temperature and added to the microcosms as described below.

### 2.2 Microcosm incubations

Microcosms containing 1 mM PO^14^C were established using combinations of labile carbon, in the form of sodium acetate or protein as bovine serum albumin [BSA]), phosphorus as phosphate, and/or nitrogen as ammonium. This concentration was selected as it is consistent with OC concentrations in Georgia coastal estuarine systems from which the microbial community inoculum was derived (400-3000 μM C) (Alberts & Takács, 1999). BSA was selected as a representative protein source due to its well-defined chemcial character and because it has frequently been used as a model protein in aquatic biogeochemical research. The labile carbon, N, and P were added at final concentrations of 500 μM-C, 75 μM-N, and 4.7 μM-P, respectively. BSA contains both C and N, at a ratio of 6.6 C:N. Thus, the concentration of inorganic N added to select microcosms was chosen to match this ratio. The P concentration was selected based on the Redfield ratio for N:P of 16. The treatments were as follows: (1) sodium acetate (250 μM acetate or 500 μM-C); (2) protein plus P (500 μM-C as BSA, 75 μM-N as BSA, 4.7 μM K_2_HPO_4_); (3) N (75 μM NH_4_Cl); (4) N plus P (75 μM NH_4_Cl, 4.7 μM-K_2_HPO_4_); and (5) control treatment with no C, N or P addition.

Microcosms were constructed as follows: the natural microbial community was obtained by pre-filtering a sample of estuarine water (from Skidaway Island, Georgia) using a Whatman GF/A filter (Whatman, GE Healthcare Bio-sciences Corporation, Piscataway, NJ; nominal pore size 1.6 μm) to reduce grazer abundance. Prefiltered estuarine water was then filtered onto a 0.22 μm filter (Supor-200 Pall Corp, Ann Arbor, MI). Cells captured on the second filter were resuspended into artificial seawater (Sigma Sea Salts, 15.0 g/L). The cell suspension was mixed and then 3.9 mL was dispensed into master mixes for each treatment with 383 mL artificial seawater (15.0 g/L, adjusted to pH 8.1) for a targeted cell density of 10^6^ cells ml^-1^. C, N and P were added to the master mixes as appropriate for each treatment, and PO^14^C (0.3779 μCi/ 1 mg PO^14^C) was added to each master mix for a final concentration of 1 mM OC. Sixty-five mL of each master mix was dispensed after gentle mixing into five replicate, 125 mL serum vials and capped with gastight butyl stoppers (National Scientific Supply, Rockwood, TN), leaving 60 ml of atmospheric headspace. The microcosms were then incubated in an incubator at 25 °C in the dark.

### 2.3 ^14^C measurements

Throughout the course of the first 36 days of incubation, samples were collected to monitor the concentrations of total ^14^C labeled organic carbon (O^14^C), particulate organic carbon (PO^14^C) and dissolved inorganic carbon (DI^14^C). Total O^14^C was measured on days 0, 1, 2, 3, 6, 8, 10, 14, 17, 20, 22, 27, 30 and 36; PO^14^C was measured on days 0, 1, 2, 8, 14, 22, 30 and 36; and DI^14^C was measured on days 1, 2, 3, 6, 8, 10, 14, 17, 20, 22, 27, 30 and 36. In all cases, 0.5 ml samples were collected from serum vials using a 22.5-guage needle and a 1 ml syringe. To quantify total O^14^C, the sample was added to a 20 ml scintillation vial preloaded with 50 μL of 10% H_2_SO_4_, to drive off ^14^CO_2_. Samples were allowed to degas for 15 minutes in a fume hood. Finally, 5 ml of Ecoscint scintillation cocktail (National Diagnostics, Mississauga, OH) was added to each serum vial. To quantify PO^14^C, the samples were filtered through a 0.22 μm filter polycarbonate filter (Millipore, Billerica, MA). The filters were added to scintillation vials containing 5 mL of scintillation fluid. To quantify DI^14^C, samples were initially stored with 50 μL of 1 M NaOH. Just prior to measurement with a Perkin-Elmer TriCarb 2910-TR scintillation counter, samples were acidified by the addition of 0.5 mL of 0.2 M HCl. CO_2_ was trapped by bubbling a stream of air through the sample into a 20 mL scintillation vial with a Teflon-septum cap containing 10 mL modified Woellers solution (50% scintillation fluid/50% β-phenylethylamine) for 20 minutes (Steen *et al*., 2012). Tests with NaH^14^CO_3_ standards indicated ^14^CO_2_ trapping efficiency was at least 95%. For all scintillation measurements, vials were vortexed, allowed to ‘rest’ for 24-72 hours, and vortexed again prior to measurement in order to minimize particle-induced quenching.

### 2.4 Modeling ^14^C data

Total O^14^C and PO^14^C data were modeled assuming a reactive fraction, which decayed according to first-order kinetics, plus an unreactive fraction, in accordance with Eq. 1:

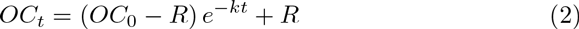

where *OC_t_* is the concentration of total OC or POC at time *t, OC*_0_ is the initial total OC or POC concentration, *R* is the concentration of recalcitrant OC or POC (modeled here as totally unreactive, in contrast with the way the term is used elsewhere in this paper), *k* is the first-order degradation rate constant, and *t* is the incubation time. These models were fit to the data using nonlinear least squares regressions, with *k* and *R* as fitted parameters and *OC*_0_ as a constant determined from measurements of the source phytoplankton (960 μM-C for total OC, 926 μM-C for POC). CO_2_ production was modeled similarly (Eq. 3), assuming that the only source of ^14^CO_2_ was the remineralization of degraded, phytoplankton-derived O^14^C.

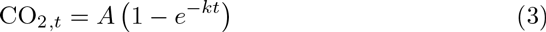

95% confidence intervals were calculated using a Monte Carlo algorithm as implemented in the `propagate` R package. For the CO_2_ data, priming at time *t* was defined as

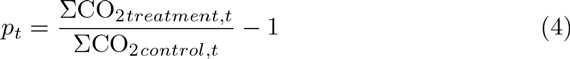

Because observed ^14^CO_2_ concentrations were non-normally distributed and temporally autocorrelated, a custom permutation test was used to test the null hypothesis that the kinetics of CO_2_ production in each treatment were different from that in the control. In this approach, which was an implementation of the generic permutation test described by Good (2013, p. 175), treatment and control labels at each timepoint were randomly shuffled, the resulting data for each reshuffled treatment were fit to Eq. 3. Priming for each permuted synthetic dataset was calculated as in Eq. 4 from the fits to Eq. 3. 95% confidence intervals for the size of the null effect on each day, including days on which ^14^CO_2_ was not measured, were calculated as the band containing 95% of priming observations out of an ensemble of 1000 randomly permuted data sets. This procedure was chosen to be insensitive to non-normality and autocorrelation, and to allow determination of whether priming occurred between measurement timepoints.

### 2.5 Potential extracellular enzyme activities

Activities of three different extracellular enzymes were assayed during the course of the incubations on days 0 (3 hours after the start of incubations), 7, 16, 21, 29 and 35. β-glucosidase was assessed using 4-methylumbelliferyl-β-D-glucopyranoside (MUB-β-glu; Sigma-Aldrich, St. Louis, MO) at a final concentration of 200 μM. Leucyl aminopeptidase was assessed using L-leucine-7-amido-4-methylcoumarin (Leu-AMC; Chem-Impex International Inc., Wood Dale, IL) at a final concentration 400 μM. Alkaline phosphatase was assessed using 4-methylumbelliferyl phosphate (MUB-PO_4_; Chem-Impex International Inc, Wood Dale, IL) at a final concentration 50 μM. At each measurement timepoint, 0.5 ml of each sample was added to 0.5 ml artificial seawater buffer and a small volume of substrate (MUB-β-glu: 20 μL, Leu-AMC: 20 μL, MUB-PO4: 50 μL). Cuvettes were capped and shaken and incubated at 22 oC. Fluorescence was periodically measured using a QuantiFluor ST single-cuvette fluorimeter over the course of approximately 2 hours as described in (Steen & Arnosti, 2013). Fluorescence values were calibrated with 4-methylumbelliferone and 7-amido-4-methylcoumarin as appropriate.

### 2.6 Cell counts

Cell densities were assessed on days 1, 3, 6, 10, 14, 17, 20, 22, 27, 30, 36 and 57 days by microscopic direct counting following (Ortmann & Suttle, 2009). 0.5 mL of sample were taken from replicate A of each treatment and stored in cryovials. 10 μL of 25% filter-sterilized glutaraldehyde was added to the samples. Samples were stored at -80*°*C. 100 μL of sample was added to 900 L of water. 50 μL of SYBR gold (25X) was added to each sample. Samples were incubated in the dark for 15 minutes. Stained samples were vacuum filtered through a 0.22 μm filter. The filter was removed and placed on a glass slide. 20 μL of antifade solution (480 μL 50% glycerol / 50% PBS; 20 μL p-phenylenediamine) was added on top of the filter on the slide before placing a cover slip on the slide. Bacteria were manually enumerated using a Leica CTR6000 microscope (Leica Microsystems, Buffalo Grove, IL).

### 2.7 Fluorescence spectroscopy of dissolved organic matter

Based on preliminary evidence that conditions in the treatments had begun to converge by 36 days, after 57 days we assessed the character of remaining DOM in selected samples using excitation-emission matrix (EEM) fluorescence spectroscopy. Due to the radioactive nature of the samples, fluorescence spectra were measured in sealed 1 cm × 1 cm methacrylate cuvettes (Fisher Scientific, Waltham, MA), which are advertised as transparent above 285 nm. In order to control for potential variability in optical properties among cuvettes, a Milli-Q water blank was measured in each cuvette prior to adding sample. For each measurement, a blank UV-vis absorbance scan was collected using Milli-Q water water on a Thermo Scientific Evolution 200 series spectrophotometer, and a blank fluorescence scan was collected on a Horiba Jobin Yvon Fluoromax 4 fluorescence spectrometer (Horiba Scientific, Kyoto, Japan). The excitation scan was from 240-450 nm in 5 nm increments, and the emission scan was from 250-550 nm in 2.5 nm increments. Finally, the Milli-Q water was removed from the cuvette, sample water was added and diluted 50% with Milli-Q water, and a sample fluorescence scan was collected using the same instrument settings. Sample 5B, which had an unacceptable blank, was discarded.

UV scans indicated that the methacrylate cuvettes began to absorb light below about 290 nm, so all excitation and emission wavelengths shorter than 295 nm were discarded. Sample fluorescence spectra were then corrected for inner-filtering effects, blank-subtracted, normalized to the appropriate days Raman spectrum, and masked for Raman and Rayleigh scattering.

BSA was the only fluorescent priming compound. For that reason, an initial fluorescence sample was taken from the control treatment prior to the addition of any priming compounds, and a separate initial sample was taken from the +BSA+P treatment to assess the fluorescence characteristics of the added BSA. Duplicate final samples were taken after 57 days incubation from each treatment.

EEMs data analysis techniques can be highly sensitive to the specific conditions under which fluorescence EEMs were measured (Cory *et al*., 2010). Since our EEMs were collected using a nonstandard cuvette type at a restricted set of wavelengths, we present the data qualitatively.

### 2.8 Data analysis

Data were analyzed using the R statistical platform (R Core Team, 2015) and visualized using the ggplot2 package (Wickham, 2009). All raw data and data-processing scripts are available at http://github.com/adsteen/priming2015.

## 3 Results

### 3.1 Character of ^14^C-labeled phtoplankton -derived OM

To generate less-reactive organic matter for microcosm studies, a culture of the marine phytoplankton species *Synechococcus* sp. CB101 was first grown in the presence of ^14^C -labeled bicarbonate. The labeled biomass was then subject to degradation by an estuarine microbial community for 45 days. At the end of the incubation period, 45 +/- 4 % of the initial phytoplankton O^14^C remained (Fig. 1) consistent with a half-life for phytoplankton OC of 36 +/- 2 days based on a first-order decay kinetics.

**Figure 1.**
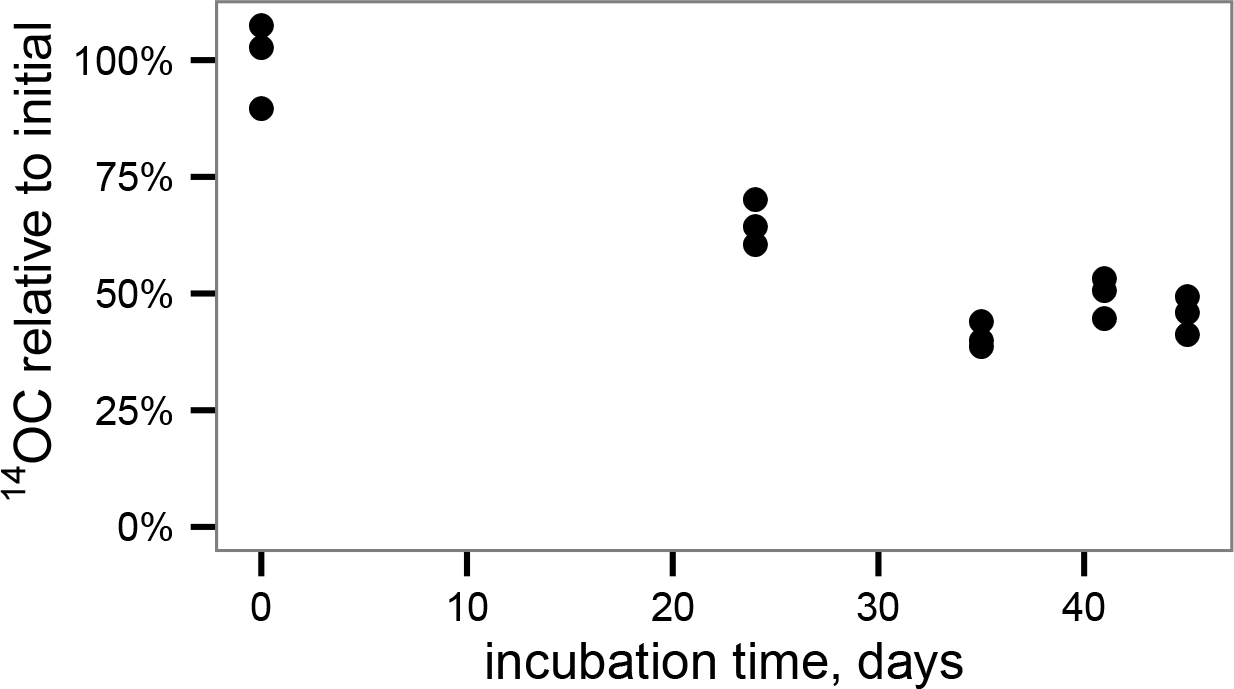
Degradation of ^14^C-labeled phytoplankton -derived OM by an estuarine microbial community yields relatively recalcitrant, ^14^C-labeled OM for use in microcosm experiments.

### 3.2 Decay of total and particulate OC

Total ^14^OC (i.e., D^14^OC+P^14^OC) and P^14^OC decayed according to similar kinetics (Fig. 2). POC in the +BSA+P treatment decayed with a faster rate constant (0.62 +/- 0.46 day-1) than any other treatment (in the range of 0.06-0.19 day^-1^, with error of 0.06-0.08 day^-1^). Substantial noise in the data obscured any other differences that might have existed in decay rate constant or concentrations of degraded, phytoplankton-derived OM.

**Figure 2.**
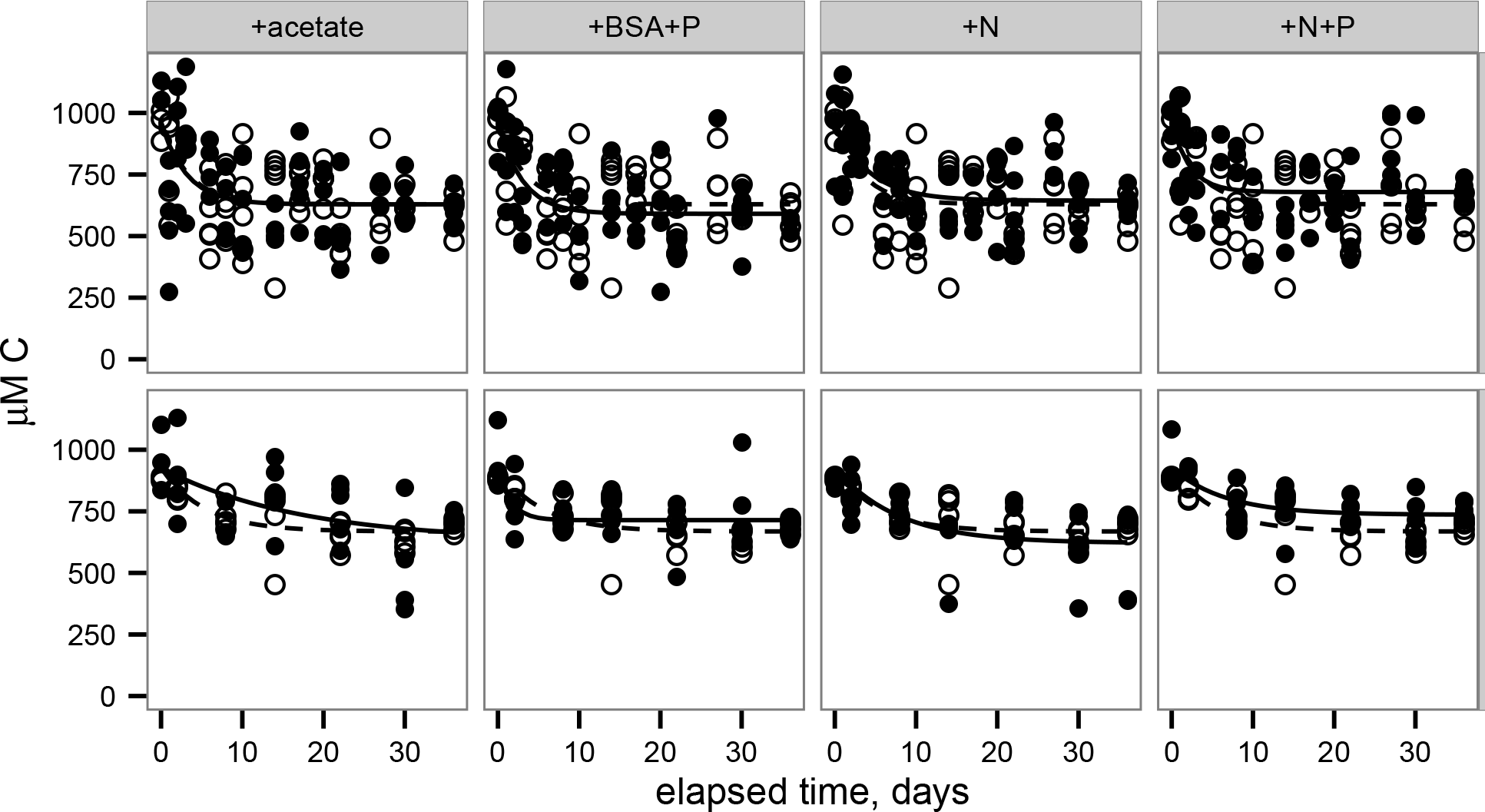
Degradation of ^14^C-labeled phytoplankton-derived OM by an estu-arine microbial community yields relatively recalcitrant, ^14^C-labeled OM for use in microcosm experiments. Remineralization of total OC (top row) and particu-late OC (bottom row). Lines indicate the nonlinear least squares regressions to Eq. 2 (provided in Methods). Filled circles and solid lines indicate data from each treatment, as indicated across the top panels. Open circles and dashed lines indicate control data and are repeated in each panel for reference.

### 3.3 CO_2_ production and priming

^14^CO_2_ production was faster in the +BSA+P treatment than in the control, indicating a positive priming effect which was distinguishable from zero (p <0.05) from day 1 through day 21 (Fig 3; Table ??). The +N+P treatment also increased the rate of ^14^CO_2_ production relative to control (Table 2). The rate constant for ^14^CO_2_ production was also larger in the +N+P treatment than the control (p <0.05) but the extent of priming in this treatment was never distinguishable from zero for an alpha of 0.05.

**Table 1.** Modeled rate constants (*k*) and modeled recalcitrant organic carbon concentration (*R*) for total O^14^C and PO^14^C in each incubation. *k* and *R* were determined according to Eq. 2 (provided in the Methods). ??

| Test | Treatment | $k$, day<sup>-1</sup> | $R$, $\mu\text{M C}$ |
| --- | --- | --- | --- |
| total | +acetate | 0.32 +/- 0.14 | 630 +/- 29 |
| total | +BSA+P | 0.33 +/- 0.11 | 590 +/- 25 |
| total | +N | 0.24 +/- 0.076 | 640 +/- 24 |
| total | +N+P | 0.44 +/- 0.2 | 680 +/- 22 |
| total | control | 0.3 +/- 0.12 | 630 +/- 24 |
| POC | +acetate | 0.058 +/- 0.059 | 630 +/- 140 |
| POC | +BSA+P | 0.62 +/- 0.46 | 710 +/- 21 |
| POC | +N | 0.13 +/- 0.063 | 620 +/- 41 |
| POC | +N+P | 0.15 +/- 0.077 | 740 +/- 23 |
| POC | control | 0.19 +/- 0.062 | 670 +/- 19 |

**Table 2.**
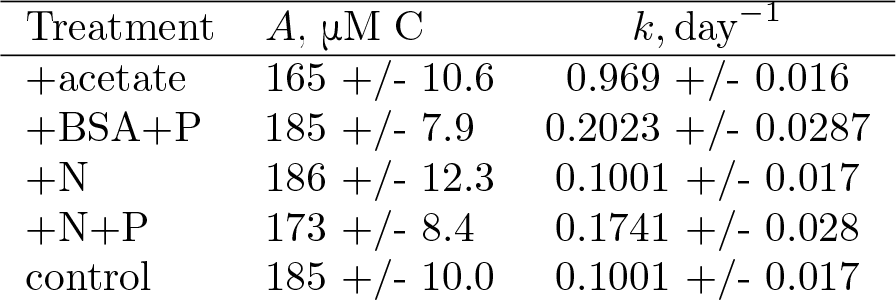
odeled asymptotes *(A)* and rate constants *(k)* for CO2 production in each incubation.

**Figure 3.**
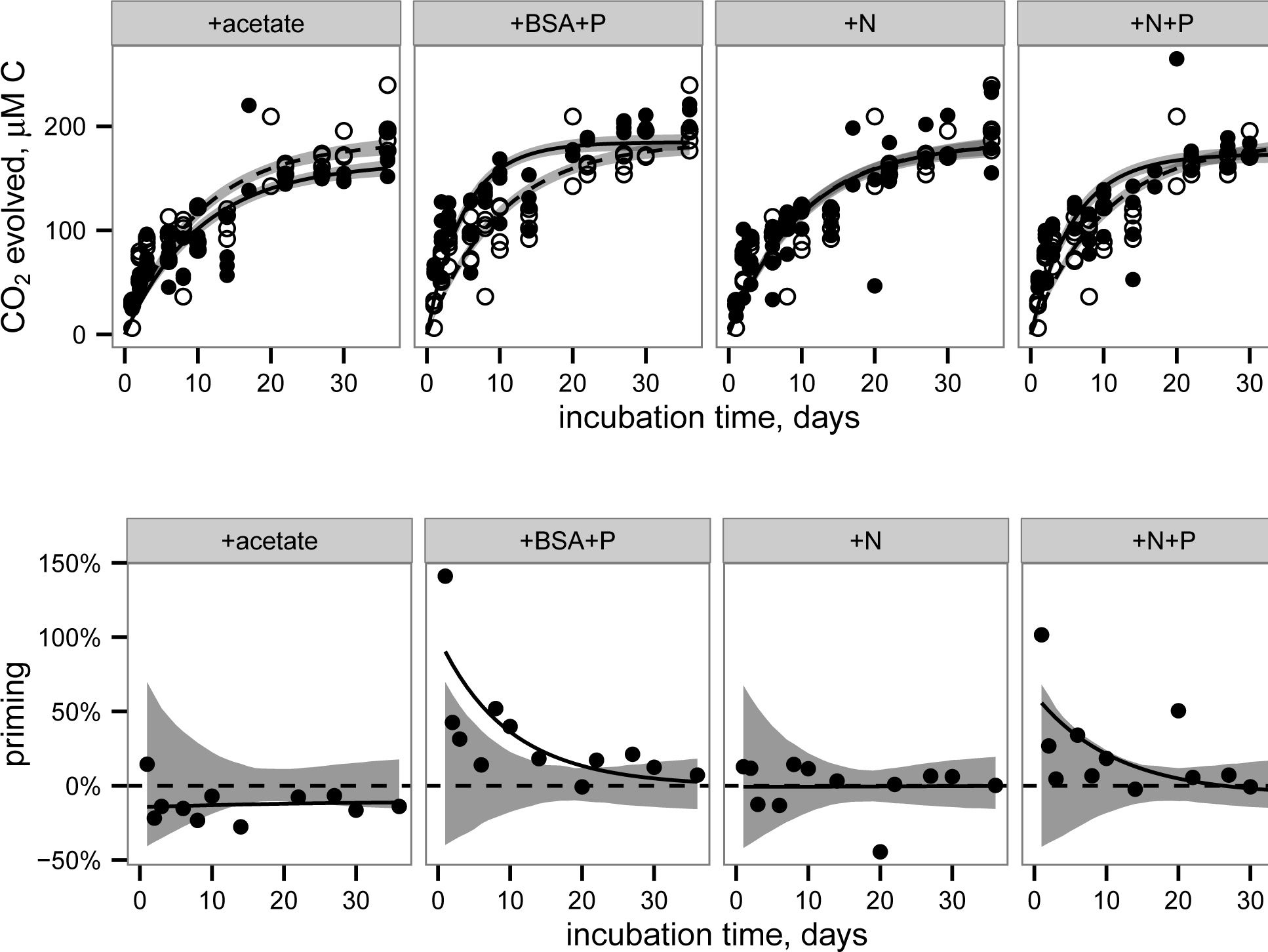
CO_2_ production (top row) and priming (bottom row) in each treatment. Top row: Filled circles and solid indicate data from treatments. Open circles and dashed lines indicate data from the control (i.e., no added compounds) and are repeated in each panel for reference. Lines indicate best fits to Eq. 3 (provided in Methods). Shaded bands indicate standard error of the model fits estimated by a Monte Carlo technique. Bottom row: Priming, calculated according to Eq. 4 (provided in Methods). Circles indicate priming calculated from the average CO2 concentrations at each timepoint. Solid lines represent priming calculated from the fit lines shown in the top panel for each corresponding treatment. Shaded bands indicate the region that is indistinguishable from zero priming.

^14^CO_2_ production in the +acetate treatment was slightly slower than in the control, consistent with a negative or anti-priming effect; this effect was significant between day 14 and day 24, and the ^14^CO_2_ production in the +N treatment was indistinguishable from the control. While the magnitude of anti-priming in the +acetate treatment was nearly constant throughout the incubation, positive priming in the +BSA+P treatment (and the +N+P treatments, if the observed priming in that treatment was not due to experimental error) was maximal at the first timepoint after labile organic matter was added, and decreased steadily thereafter. After 30-36 days of incubation, the total amount of ^14^CO_2_ remineralized was indistinguishable among all treatments.

After 36 days of incubation, our quantification indicated that more TOC was removed from the system (320-370 μM) than CO_2_ was produced (165-186 μM). The average deficit of 147 ± 30 μM likely represents biofilms attached to the incubation vessel walls, which would have been missed by our sampling method.

### 3.4 Cell abundance and extracellular enzymes

Cell abundances in the incubations increased from approx. 1.0 × 10^6^ cells ml^-1^ in each treatment after 1 day of incubation to 1.4-2.5 × 10 ^6^ cells ml ^-1^ after 57 days of incubation, with relatively little difference among treatments (Fig 4). However, substantial differences among treatments occurred during the course of the incubation. In the +BSA+P treatment, cell densities quickly increased to a maximum of 1.2 × 10^7^ cells ml^-1^ after 3 days and then decreased steadily through the end of the incubation. Other treatments were characterized by an initial peak at 6 days incubation. Cell abundance in the +N treatment remained roughly constant after 6 days, whereas the control, +acetate, and +N+P treatments, followed by a minimum in cell abundance at approximately 17 days, and, in the case of the +acetate treatment, a second, larger peak in cell abundance at 27 days.

**Figure 4.**
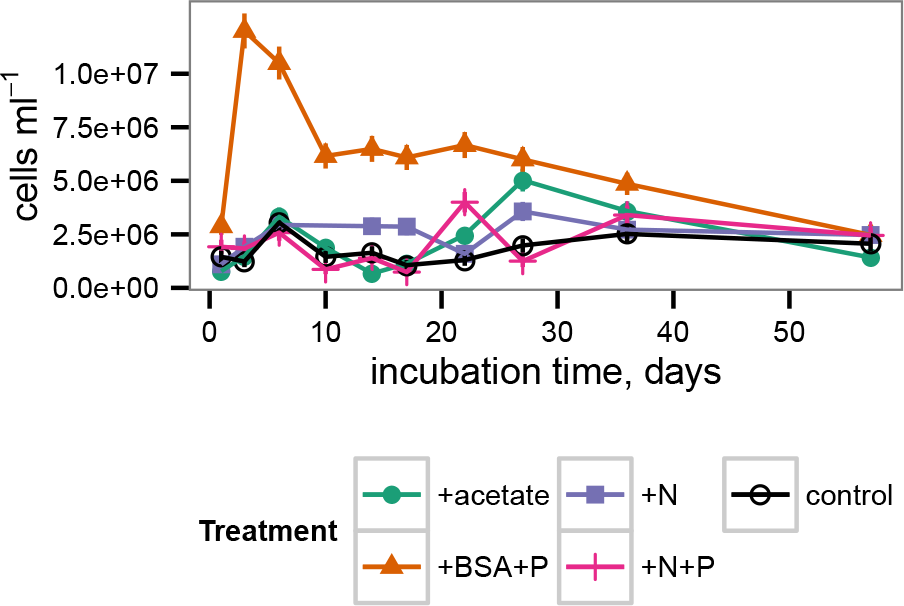
Cell abundance during the incubation. Error bars represent standard error of cell counts.

Potential activities of extracellular enzymes also varied as a function of both time and treatment (Fig 5). β-glucosidase activities were generally indistinguishable from zero throughout the incubation for all treatments. Leucyl aminopep-tidase activities were far greater in the +BSA+P treatment than in any other treatment, although activities were significantly greater than zero in each treatment. The timecourse of leucyl aminopeptidase activities followed cell counts closely. Alkaline phosphatase activities were also greater in the +BSA+P treatment than in any other treatment, but the timecourse of activities followed a different path than the timecourse of cell counts: the maximum value was at 17 days rather than 6 days, and the peak in activities was less dramatic than either the peak in leucyl aminopeptidase activities or cell counts. While most measures of biological activity ceased at day 35 due to limited sample volume, a final measurement of cell density was made at day 57 and found to range from 1.4 × 10^6^ cells ml^-1^ (+acetate treatment) to 2.5 × 10^6^ cells ml^-1^ (+BSA+P, +N, +N+P treatments).

**Figure 5.**
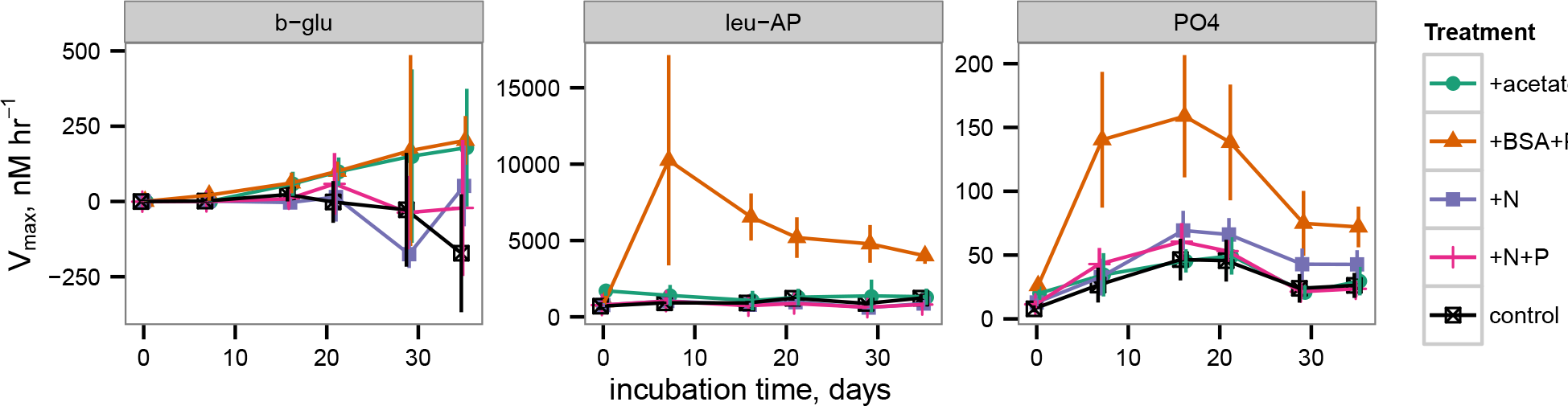
Potential extracellular enzyme activities during the incubation. bglu represents β-glucosidase, leu-AP represents leucyl aminopeptidase, and PO4 represents alkaline phosphatase.

### 3.5 Chemical transformations of DOM

At the conclusion of the incubation period (day 57), the remaining sample volume was sacrificed for excitation-emission matrix (EEM) fluorescence spectro-scopic analysis and compared with samples preserved from the first day of the incubation. The intensity of the FDOM signal increased in all samples over the course of the incubation (Fig 6). The nature of the signal, as revealed by EEM, however, did not vary much by treatment, with the exception of the +BSA+P treatment. In this treatment, the protein peak from the added BSA (visible at the bottom of the panel for the initial +BSA+P treatment in Fig 6) dominated the phytoplankton degraded, phytoplankton-derived OM signal. By the end of the incubation, however, there was no distinct protein signal, and the overall form of the EEM in the +BSA+P treatment was considerably more intense but similarly shaped to the signals from the other treatments.

**Figure 6.**
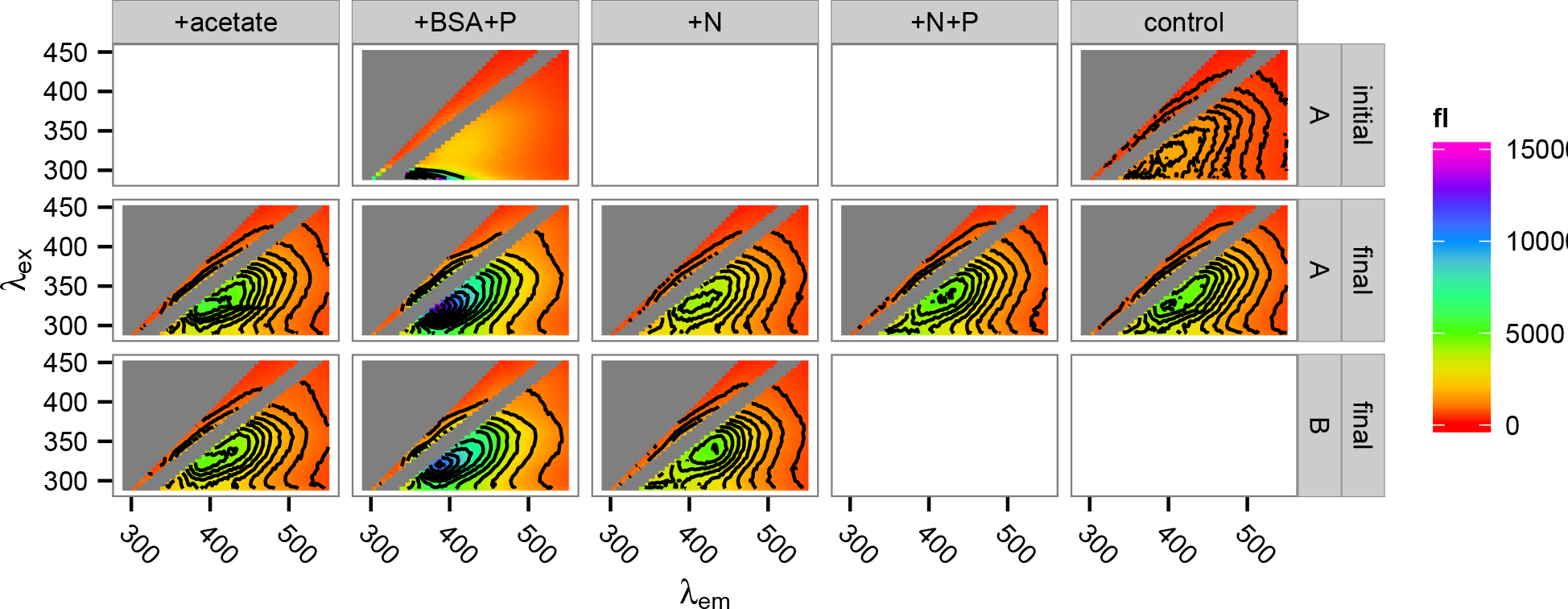
Fluorescence spectra of incubation DOM at the start and at the end of the incubations. Top row: spectra of the +BSA+P treatment and the control treatment at time zero (‘initial’). Middle and bottom rows: replicate spectra after 57 days incubation (‘final’). Insufficient sample remained for duplicate measurement of the +N+P and control samples after 57 days. ‘A’ and ‘B’ in the right-side panel labeled refer to incubation replicates.

## 4 Discussion

### 4.1 Reactivity of source O^14^C

For this study, we selected a representative strain of phytoplankton, *Synechococcus* sp. CB0101, which was originally isolated from the Chesapeake Bay (Marsan *et al*., 2014). *Synechococcus* can account for a substantial fraction of total phototrophic cells, chlorophyll a, and primary production in estuaries (Ning *et al*., 2000; Pan *et al*., 2007; Wang *et al*., 2011). During preparation, the phytoplankton-derived organic matter used in this experiment decayed with a half-life of 36 ± 2 days, consistent with semi-labile estuarine DOC (Raymond & Bauer, 2000). Although the phytoplankton-OM decay data here were too sparse to accurately model with a multi-G model (Fig. 1), the success of more complex diagenetic models indicates that organic matter becomes less reactive as it is oxidized by microorganisms (e.g., Røy *et al*., 2012). It is therefore likely that the remaining organic matter at the end of the pre-degradation phase was less reactive than the halflife of 36 days would suggest.

### 4.2 Priming as a transient effect

Recalcitrant OM was remineralized up to 100% faster in the +BSA+P treatment than in the control, but this effect was transient (Fig 3). After about 30 days, roughly the same amount of recalcitrant OM had been mineralized in each experiment. Cell densities (Fig 4) and enzyme activities (Fig 5) also converged towards the end of the experiment. Fluorescence spectroscopy indicated that, after 57 days of incubation, the composition of fluorescent DOM was indistinguishable among all treatments except for +BSA+P. In that treatment a large protein-like peak persisted at the end of the incubation. Other than the large protein-like peak, post-incubation fluorescence spectra of the +BSA+P treatment were qualitatively similar to post-incubation spectra for the other treatments (Fig 6).

Interestingly, Catalán *et al*. (2015) recently found no evidence of priming in Swedish lakes. That study contained a very large number of experimental treatments, but only a single timepoint, after 35 days of incubation, whereas in the experiment reported here, priming effects were no longer observable after 21 - 24 days. The priming effect arises from interactions between disparate microorganisms and pools of organic carbon and nutrients (Blagodatskaya & Kuzyakov, 2008). Given the complexities of these interactions, it is likely that the magnitude, direction and timing of priming varies substantially among aquatic environments.

### 4.3 Priming versus stoichiometric control on CO_2_ production

The results provide evidence of faster OM mineralization in the presence of added protein plus phosphate (+BSA+P treatment) and possibly added inorganic N and phosphate (+N+P), but not inorganic N alone (+N). These data suggest that heterotrophic metabolism of recalcitrant OM was limited in part by phosphorus. It is important to note that the factors limiting the remineral-ization of recalcitrant OM may differ from the factors limiting overall bacterial production. Because this experiment involves comparing treatments that received additional nutrient inputs to a control in which no nutrients were added, it is important to distinguish potential stoichiometric effects of nutrient addition from a priming effect. The addition of N and P in the +BSA+P, +N+P, and +N treatments could be expected to spur remineralization of excess ^14^CO_2_ relative to the control, purely to maintain stoichiometric balance. However, two lines of evidence indicate that some fraction of the excess ^14^CO_2_ observed in the +BSA+P treatment was due to priming by BSA. First, First, the magnitude of the effect in the +BSA+P treatment was roughly twice as large as in the +N+P treatment, despite the identical N:P stoichiometry in the two treatments. Second, the fact that the effects observed here were transient is difficult to reconcile with stoichiometric effects: we are not aware of a mechanism by which stoichio-metric effects could cause the ^14^CO_2_ production in the control to ‘catch up’ to that in the experimental treatments, as we observed here, without additional input of nutrients, whereas priming effects are well-known to be time-dependent (Blagodatskaya & Kuzyakov, 2008).

### 4.4 Potential mechanism of priming

Cell abundances, leucyl aminopeptidase activity and phosphatase activity all increased substantially and rapidly in the +BSA+P treatment (Figs 5 and 6). This is consistent with a scenario in which cells grew rapidly using BSA as a substrate, producing excess leucyl aminopeptidase, which released bioavailable compounds (e.g. amino acids) from the protein-like organic matter that comprises the major fraction of organic N in degraded organic matter (Nunn *et al*., 2010; McCarthy *et al*., 1997). Kuzyakov et al Kuzyakov *et al*. (2000) cite changes in microbial biomass as a primary mechanism of priming in soils. Surprisingly, alkaline phosphatase activity also increased in the +BSA+P treatment, despite the substantial addition of P in that treatment. Some marine bacteria produce alkaline phosphatase constituitively (Hassan & Pratt, 1977), which may account for the observed increase in alkaline phosphatase activity in the +BSA+P treatment here. Alternatively, since the peak in phosphatase activity occurred at 17 days while cell abundance was declining, it is possible that the extracellular phosphatase enzymes may have been released from cytoplasm as cells lysed following the peak in cell abundance at day 3. Alkaline phosphatase can cleave phosphate from phosphoproteins (Mellgren *et al*., 1977), so the extra peptidases expressed in the +BSA+P treatment may have liberated phosphosphoproteins which induced expression of alkaline phosphatase-like enzymes. In any case, the observed increase in the activity of phosphatase provides mechanistic support for the hypothesis that addition of one compound can spur hydrolysis of chemically unrelated compounds, thereby making them bioavailable. Many aquatic extracellular peptidases (protein-degrading enzymes) are relatively promiscuous (Steen *et al*., 2015) which suggests that peptidases produced in order to degrade BSA likely hydrolyzed some fraction of the recalcitrant O^14^C as well.

The microcosms used in this study contained planktonic cells, suspended particles and flocs, and probably biofilms attached to incubation vessel walls. The physiological state of bacteria growing attached to surfaces is dramatically different than when they are unattached (reviewed in Costerton et al Costerton *et al*. (1995)). and is, therefore, an important consideration for microbial transformation studies. It is possible that the mechanisms and extent of priming differed among these microenvironments, as suggested by Catalán et al Catalán *et al*. (2015).

### 4.5 Relevance to carbon processing in estuaries

Priming in this study was substantial but transient. The relevant priming timescale observed here of days-to-tens-of-days, coincides with typical hydro-logic residence times of passive-margin estuaries (Alber & Sheldon, 1999). Priming in estuaries may therefore influence whether OC is remineralized in situ or exported to the coastal ocean.

Is the priming effect that we observed in microcosm incubations with defined substrate additions relevant to natural systems? In estuaries, degraded OM (e.g. terrestrial OM or dissolved remnants of coastal phytoplankton blooms) can come into contact with fresh DOC produced in situ (Raymond & Bauer, 2001). Marsh grasses exude substantial amounts of labile compounds, including acetate (Hines *et al*., 1994; Jones, 1998), and phytoplankton growing in estuaries likely also serve as a source of labile OM (Carlson & Hansell, 2015). Here, we have shown that estuarine microbial communities are capable of being ‘primed’ (or ‘anti-primed’) by the addition of labile OM and nutrients to mineralize recalcitrant OM more quickly. Therefore, we hypothesize that inputs of labile OM and nutrients to estuaries may influence fluxes of organic carbon between estuaries and the coastal ocean. Given the numerous environmental variables that cannot be accounted for in lab-scale experiments, this hypothesis should be tested with field-scale experiments.

## Author Contributions

ADS and AB designed the experiment. ADS, LNMQ and AB performed the experiment, analyzed the data, and wrote the manuscript.

## Acknowledgments

We thank Aron Stubbins (Skidaway Institute of Oceanography) and Mike Piehler (University of North Carolina Institute of Marine Sciences) for help collecting microbial inocula. Steven Wilhelm (UT-Microbiology) shared lab space for experiments, Annette Engel shared the fluorimeter used to collect DOM fluorescence spectra, and Kathleen Brannen-Donnelly gave valuable help with data processing.

*Funding:* This work was supported by NSF grant OCE-1357242 to ADS and AB.

